# A validated workflow for rapid taxonomic assignment and monitoring of a national fauna of bees (Apiformes) using high throughput barcoding

**DOI:** 10.1101/575308

**Authors:** Thomas J. Creedy, Hannah Norman, Cuong Q. Tang, Kai Qing Chin, Carmelo Andujar, Paula Arribas, Rory O’Connor, Claire Carvell, David G. Notton, Alfried P. Vogler

**Affiliations:** Department of Life Sciences, Natural History Museum, Cromwell Rd, London, SW7 5BD, UK; Department of Life Sciences, Silwood Park Campus, Imperial College London, Ascot, SL5 7PY, UK; Science and Solutions for a Changing Planet DTP, Department of Life Sciences, Silwood Park Campus, Imperial College London, Ascot, SL5 7PY, UK; NatureMetrics Ltd, CABI Site, Bakeham Lane, Egham, Surrey, TW20 9TY, UK; Island Ecology and Evolution Research Group (IPNA-CSIC), Astrofísico Fco. Sánchez 3, 38206 La Laguna, Tenerife, Spain; Faculty of Biological Sciences, University of Leeds, Leeds, UK, LS2 9JT; School of Agriculture, Policy and Development, University of Reading, Whiteknights, PO Box 237, Reading, UK, RG6 6AR; NERC Centre for Ecology & Hydrology, Benson Lane, Crowmarsh Gifford, Wallingford OX10 8BB, UK

**Keywords:** Pollinators, community barcoding, contamination, lllumina sequencing, double dual tagging

## Abstract

Improved taxonomic methods are needed to quantify declining populations of insect pollinators. This study devises a high-throughput DNA barcoding protocol for a regional fauna (United Kingdom) of bees (Apiformes), consisting of reference library construction, a proof-of-concept monitoring scheme, and the deep barcoding of individuals to assess potential artefacts and organismal associations. A reference database of Cytochrome Oxidase subunit 1 (*cox1*) sequences including 92.4% of 278 bee species known from the UK showed high congruence with morphological taxon concepts, but molecular species delimitations resulted in numerous split and (fewer) lumped entities within the Linnaean species. Double tagging permitted deep illumina sequencing of 762 separate individuals of bees from a UK-wide survey. Extracting the target barcode from the amplicon mix required a new protocol employing read abundance and phylogenetic position, which revealed 180 molecular entities of Apiformes identifiable to species. An additional 72 entities were ascribed to mitochondrial pseudogenes based on patterns of read abundance and phylogenetic relatedness to the reference set. Clustering of reads revealed a range of secondary Operational Taxonomic Units (OTUs) in almost all samples, resulting from traces of insect species caught in the same traps, organisms associated with the insects including a known mite parasite of bees, and the common detection of human DNA, besides evidence for low-level cross-contamination in pan traps and laboratory steps. Custom scripts were generated to conduct critical steps of the bioinformatics protocol. The resources built here will greatly aid DNA-based monitoring to inform management and conservation policies for the protection of pollinators.

## INTRODUCTION

Widespread declines in pollinator populations are raising the alarm about the future of global biodiversity and agricultural productivity (Garibaldi *et al*. 2013; Hallmann *et al*. 2017; Lever *et al*. 2014), driven by the combined effects of habitat loss, introduction of non-native and invasive species, pathogens and parasites, and various other factors contributing to environmental change (Vanbergen *et al*. 2013). Landscape effects on pollination of crops through agricultural intensification, particularly those of monoculture crops, have led to significant changes in pollinator communities (Kennedy *et al*. 2013; Ricketts *et al*. 2008), with obvious economic implications for the agricultural sector and pollination services worth hundreds of millions of pounds in the United Kingdom alone (Potts *et al*. 2010). However, these trends in species distribution and abundance are difficult to quantify, unless solid methodologies for monitoring at regional levels can be implemented. Thus there is an urgent need to develop strategies for large-scale and long-term systematic monitoring of pollinator populations, to better understand the impacts of declines on pollination services to crops and wild plants, and inform policy decisions and conservation efforts.

Current evidence of change in pollinator populations in the United Kingdom comes primarily from analyses of records of species occurrence submitted by volunteer recorders (e.g. (Biesmeijer *et al*. 2006). While these allow for the analysis of large-scale changes in species distributions, they provide no information on abundance or local population size, and are known to be temporally and spatially biased (Isaac & Pocock 2015). Instead, pan traps have been proposed as the most effective method for systematic monitoring of bee diversity in European agricultural and grassland habitats (Westphal *et al*. 2008). Species identification is usually performed with morphological analysis by expert taxonomists, but there is a growing need for alternative methods, in particular because the great species diversity and large quantity of specimens from mass trapping make them challenging and costly to identify (Lebuhn *et al*. 2013).

This study applies high throughput sequencing (HTS) techniques to assess bee diversity and abundance from mass-trapped samples, using a rapid DNA barcoding approach suitable to individually assay the thousands of specimens potentially generated in the course of a large-scale monitoring scheme. The first step in this process was the generation of a well curated reference database that links each DNA sequence to a species name, using the Cytochrome *c* Oxidase subunit I *(cox1)* barcode marker (Hebert *et al*. 2003), which provides good species discrimination in Hymenoptera (Smith *et al*. 2008). The more recent approach of ‘metabarcoding’, by which entire trap catches are subjected to amplicon sequencing in bulk, produces species incidence data based on the mixed sequence read profile (Yoccoz *et al*. 2012). Current HTS protocols can maintain the individual information of thousands of samples using unique tags in the initial PCR prior to pooling for illumina sequencing with secondary tags, which allows sequences to be traced back to the associated specimen (Arribas *et al*. 2016; Shokralla *et al*. 2015). The great sequencing depth of this high-throughput barcoding (HT barcoding) methodology may also reveal DNA from organisms internally or externally associated with a target specimen or as a carry-over from other specimens in the trap.

The current study on the regional-scale pollinator fauna, focused on the bees (Hymenoptera: Apiformes) of the United Kingdom, illustrates the required steps from generating a barcode reference database for the 278 species of bees known from the UK, which was then used for the identification of samples gathered as part of a pilot study for a national monitoring scheme based on short barcode sequences obtained with HTS. Agreement between morphological and molecular identifications was assessed. In addition, the deep-sequencing approach allowed the assessment of organisms associated with the target specimens, as well as cross-contaminations from other species present in the traps or from specimen handling and laboratory procedures.

## MATERIALS AND METHODS

### Building a regional reference database

A *cox1* reference database was generated from DNA barcoding of specimens of bee species known to occur in the UK according to the list of Falk and Lewington (2015) and notes from various sources maintained by co-author DGN. Most specimens were caught by hand netting and identified by DGN, using the latest keys available at the time (Amiet *et al*. 2001, 2004, 2010; Amiet *et al*. 2007; Amiet *et al*. 2014; Bogusch & Straka 2012; Falk & Lewington 2015) (Benton 2006; Mueller 2016). Identifications had to draw on these various references because the comprehensive key of Falk & Lewington (2015) became available only part way through the study, while some identifications were also cross-checked between different publications. Specimen data for morphological vouchers are available at the Natural History Museum Data Portal (data.nhm.ac.uk) http://dx.doi.org/10.5519/0002965. Sequences are available at BOLD (Barcode-of-Life Datasystem) under the BEEEE project label. Additional specimens were obtained using pan traps from the survey described below. The reference set included all available unique UK species as determined by morphology, with multiple specimens per species where available. These within-species replicates allowed inclusion of specimens from across the geographical range of widely-distributed species, identified by different taxonomists and/or belonging to known species complexes.

DNA was extracted from a single hind leg using a Qiagen DNeasy Blood and Tissue Kit, after the specimens were incubated at 56°C in the extraction buffer (ATL and Proteinase K) overnight in a shaking incubator at 75 rpm. The complete ‘barcode region’ (658 bp) of *cox1* was amplified using newly designed primers (BEEf TWYTCWACWAAYCATAAAGATATTGG and BEEr TAWACTTCWGGRTGWCCAAAAAATCA), based on an alignment of 84 mitochondrial genomes from 22 genera. PCR and sequencing using ABI dye terminator sequencing followed standard procedures (Supplementary Material). The sequences were added to BOLD in the project BEEEE, along with Syrphidae barcodes that were sequenced at the same time.

Sequences were aligned using the *MAFFT* v1.3. (Katoh *et al*. 2009) plugin in *Geneious*. Alignments were used for molecular distance-based and coalescence-based species delimitation. (1) BOLD BINs (Barcode Identification Numbers) were automatically generated for sequences that are uploaded onto its database (Ratnasingham & Hebert 2013). These BINs are formed using a refined single linkage network, which combines sequence similarity metrics and graph theory. (2) The GMYC (Generalized Mixed Yule Coalescent) method determines species limits based on a shift in the rate of branching along the root-to-tip axis of the phylogenetic tree, separating the speciation (Yule) processes from fast branching rate expected within population under a neutral coalescent (Fujisawa & Barraclough 2013). The analysis was done on phylogenetic trees constructed separately for each genus using BEAST 1.8.1 (Drummond & Rambaut 2007), which was used as the input for the GMYC analysis (Tang *et al*. 2014). The GMYC was applied to genera with only a single British representative (*Apis, Anthidium, Ceratina, Dasypoda, Macropis* and *Rophites*) by adding supplementary sequences from BOLD.

New barcode sequences for species that were already in BOLD were assessed to examine whether these new sequences represented new *cox 1* haplotypes for these species. The new barcodes were searched against a set of 1754 sequences downloaded from BOLD for species known to exist in the UK using BLASTn on default parameters. Any sequences not 100% identical over the entire length of the query or subject were designated as novel haplotypes.

### Generating a test dataset from field caught samples using HTS

The reference database was used for identification of specimens obtained through the National Pollinator and Pollination Monitoring Framework (NPPMF) (Carvell et al., 2016). Mixed samples were collected with pan traps consisting of sets of water-filled bowls (painted UV-yellow, white and blue, after Westphal *et al*. 2008) from 14 sites across the UK, and further specimens were collected by netting along standardised transects running 200m from each set of pan traps (Figure 1A, Table S1; see Carvell *et al*. 2016 and supplementary materials for a full description of the sampling protocol). Bees (Apiformes) were separated from other taxa in the field, stored in 99% ethanol to preserve DNA for analysis, and transferred to −20°C as soon as possible after collection. Specimens were identified morphologically by expert taxonomists offering commercial identification services. In total, 762 bee specimens were processed (480 bees were extracted from the pan traps, and 282 specimens from the transects and further hand collecting). All specimens were stored in 99% ethanol and deposited as voucher specimens in the Molecular Collection Facility at the NHMUK.

**Fig. 1.**
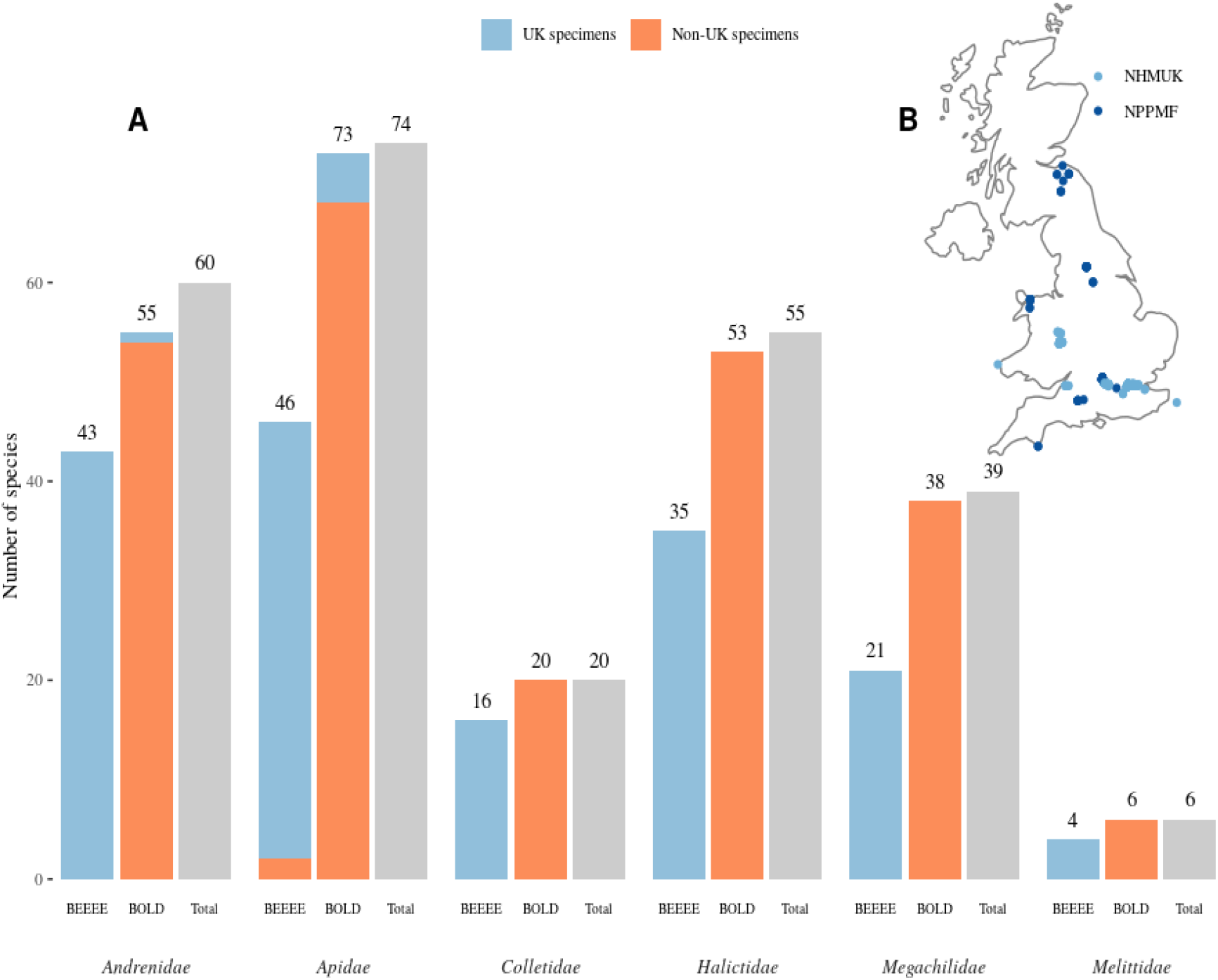
Specimens and species used in this study. A. The number of bee species of each family, dataset and geographical source from which sequences were compiled to form the reference collection, Column colours denote whether species from each dataset comprised any UK specimens, and numbers above bars give totals, The BEEEE columns denote the species sequenced as part of this study (165), which were compiled with existing BOLD sequences (245 species) to form the total number of species sequenced per family. This dataset comprises 255 of the 278 bee species in the UK. B. Sampling localities of bee specimens collected by NHMUK and the NPPMF that formed the BEEEE reference set of specimens

DNA was extracted from individual specimens by piercing the abdomen and submerging the whole specimen in lysis solution consisting 180ul ATL buffer and 20ul Proteinase K for 12 hours on a 56°C shaking incubator. DNA extractions were performed using either the Qiagen BioSprint 96 DNA Blood Kit or DNeasy Blood and Tissue kits applied to the lysate. Each DNA extract was PCR amplified for a 418bp portion of the *cox1* barcode region (Andujar *et al*. 2018). Each individual amplicon was tagged using a ‘double dual’ PCR protocol (Shokralla *et al*. 2015) to generate unique tag combinations for each bee specimen, following the procedures of Arribas *et al*. (2016). Tags were added in the initial PCR by amplification using tagged *cox1* primers employing different 6 bp sequence combinations designed with a Hamming distance of 3, with a total of 13 different tagged primer sets. In all reactions, forward and reverse primers used the same tag, so that the products of tag jumping could be removed (Schnell *et al*. 2015). Amplicons generated with different primer tags were pooled, and each pool was cleaned using Agencourt AMPure XP beads (Beckman Coulter, Wycombe, UK), prior to secondary amplification of each pool with the i5 and i7 Nextera XT indices with 96 unique MID combinations (illumina, CA, USA) and sequencing on illumina MiSeq v.3 (2×300 bp paired-end).

Perl scripts of the custom NAPtime pipeline (www.github.com/tjcreedy/NAPtime) were used to wrap bioinformatics filtering of the raw data. The 96 libraries were demultiplexed based on XT MIDs using illumina software and were further demultiplexed using NAPdemux based on the unique tags of the first-round PCR primers. This script wraps cutadapt (Martin 2011) for large demultiplexing runs, and used the default 10% permitted mismatch to the adapter sequences (permitting no errors in the 6 bp tag used) before binning reads according to their tags. Mate pairs with only one read matching the correct tag were discarded. Read quality was reviewed using FASTQC (Andrews 2010). Following demultiplexing, the NAPmerge script was used to generate a set of full-length reads for further analysis. The script invokes cutadapt (Martin 2011), PEAR (Zhang *et al*. 2014) and *USEARCH - fastq_fiiter* (Edgar 2010) to bulk remove primer sequences, assemble read pairs, and perform quality filtering. Any reads not containing a correct primer sequence, and their mates, were discarded, and any merged reads with 1 or more expected errors were removed with *fastq_filter*. This process generated a pool of complete *cox1* amplicon sequences for each of the specimens.

### Testing the utility of the reference dataset

Three methods were used to designate a single putative “high-throughput barcode” (HT barcode) sequence representing the *cox1* gene of each specimen from the set of reads. Firstly, we employed a standard metabarcoding pipeline, implemented in the NAPcluster script, to generate OTU (Operational Taxonomic Units) clusters and centroid sequences using *USEARCH*. The script includes functions from the *USEARCH* suite (Edgar 2010), starting with the data output from NAPmerge (merged and quality-filtered amplicons), and comprises the following steps: (i) filtering sequences by length; (ii) dereplication and filtering by number of reads per unique sequence, to retain only sequences represented by a set minimum of reads; (iii) denoising using the *UNOISE* algorithm (Edgar *et al*. 2011); (iv) clustering of sequences according to cluster radius and generation of an output set of OTU consensus sequences, and (v) mapping of reads to OTU clusters (using *USEARCH usearch_global* and a custom, uc parser) and generation of an output table of OTU read numbers by sample. All sequences differing from 418 bp and with only 1 copy were removed in steps (i) and (ii), and *USEARCH cluster_otus* was employed for clustering with a dissimilarity threshold of 3%. The centroid of the most abundant OTU was used as the specimen barcode.

As sequence variants may drive up the read count of an OTU and thus obscure the haplotype of the target specimen, the second method for selecting the HT barcode sequence simply chooses the most frequent read for each library, under the assumption that error-free reads represent the most abundant template DNA and thus the target specimen. Other sequences that represent nuclear mitochondrial pseudogenes (NUMTs), gut contents, internalised parasites, and cross-contamination, or result from PCR or sequencing errors, are expected be present in lower numbers. The extraction of the most frequent read was performed using a custom perl script.

The third method employed a purpose-built tool, NAPselect, that finds the most highly represented sequence among reads in an amplicon mix, as above, but then statistically and taxonomically validates this selection. The process starts with steps similar to metabarcoding: firstly, filter the batch of sequence reads by length (in this case, rejecting any sequence not 418 bp), and group sequences by identity (i.e. dereplication) recording the abundance of reads representing each unique sequence. Starting with the most abundant, unique sequences are assessed one-by-one, using bootstrapping to validate the significance of the difference in read abundance, and using BLAST to assign the sequence taxonomy. Based on the total number of reads in the sample and the number of unique sequences, the probability of a sequence occurring as frequently as the most abundant sequence by chance alone is determined using 10,000 bootstrap iterations. A p-value of 0 designates a sequence as significantly more frequent with high confidence, and less than 0.5 for low confidence, above which the entire sample is disregarded because a putative barcode sequence for the target specimen is not clearly defined. The most abundant sequence is then subjected to a BLASTn search against a local copy of the NCBI *nt* database and the hits assessed for presence in the focal taxon (in this case, Hymenoptera). Only those sequences passing both these tests are selected.

The success of these three methods, and the accuracy of the sequences they output, was tested by identifying the HT barcode sequences using the BEEEE reference collection of sequences obtained in section 1. Each of the three putative HT barcodes for each specimen were searched for matches in the BEEEE reference collection using BLASTn with default parameters. Only matches with >95% identity and overlap with the reference sequences of >400 bp were retained, and the match with the similarity was selected, using bitscore to break ties. The identity of this hit was compared against the known morphological ID for that specimen at the genus and species level. For each HT barcode selection method and taxonomic level, the number of correct molecular identifications was tallied and a proportion of failure calculated.

### Exploration of concomitant DNA in the testing dataset

The OTUs generated with the NAPcluster script (see above) allowed the exploration of co-amplified DNA from each bee specimen other than the primary *cox1* sequence. For each sample, the OTUs that did not match the NAPselect HT barcode sequence for the target specimen were designated as “secondary OTUs”. We used the OTUs for this analysis, rather than the reads, to reduce unnecessary complexity in the dataset. These OTUs were searched against a local copy of the NCBI *nt* database using BLASTn, followed by taxonomic binning using MEGAN6 Community Edition with the weighted Lowest Common Ancestor algorithm (Huson *et al*. 2016). Any OTUs assigned to Apiformes were additionally identified using BLASTn against the BEEEE reference collection and theNAPselect HT barcodes (above, section 3). In both cases, BLASTn employed default parameters, and sequences were identified as the hit with the highest identity where identity was >95% and overlap was >400 bp, with bitscore breaking ties.

NUMTs may appear as separate OTUs in metabarcode data and add spurious OTUs to the clusters derived from the true mitochondrial copy. A tree-based filtering pipeline was used to identify NUMT-derived OTUs based on the assumption that they are closely related to the corresponding mitochondrial copy, and are coincident across sequenced samples, while their copy number is lower. Thus, OTUs were considered derived from pseudogenes if they were completely coincident across samples with another closely related OTU that did match a BEEEE reference or specimen barcode, and the number of reads was significantly lower in comparison.

The resulting datasets were reconfigured for various statistics and to perform downstream calculations using R (Team 2018) packages *plyr* and *reshape2*. The OTU x sample dataset was rarefied to 400 reads per sample to facilitate valid comparison between samples using the R package *vegan* (Oksanen *et al*. 2018).

Cross-contamination among samples was tested by assessing the distribution of secondary OTUs in each sample obtained from pan trapping. Only secondary OTUs that matched a (NAPselect) HT barcode (section 2) from *another* sample were used in this analysis. Three sources of cross-contamination were considered: contamination from other individuals in the same trap, DNA mixing between specimens with the same PCR tag on a single plate, and DNA mixing between specimens with the same Nextera XT tag in a single well. For each source or combination of sources, the total possible selected barcodes were counted, and then the proportion of those that were present as secondary OTUs in a sample was calculated. For example, each well in the library preparation contained 13 specimens tagged with different sequence identifiers: if in a set of these 13 each is a different species (different HT barcodes), there are 12 possible well contaminants for any one of these samples, and so a sample containing 3 secondary OTUs from these 12 specimen barcodes would have a contamination rate of 3/12 = 0.25 from well-level contamination. As a control, the rate of contamination from all possible sources together was also scored, i.e. the proportion of secondary OTUs in a sample that matched *any* HT barcodes, out of the total number of unique HT barcodes.

One-sample *t* tests were used to assess if the mean contamination rate for each source or source combination was significantly greater than zero. To compare between sources against the control, the effect of source on contamination rate was fitted in a quasi-binomial ANOVA, setting the control as the reference level.

## RESULTS

### A reference database of UK bees

A total of 355 bee specimens were newly sequenced for the COI barcode to generate the reference set, representing 165 Linnaean species. These new sequences were added to 1754 full-length barcode sequences downloaded from the Barcode of Life Database (BOLD) for a total of 2109 sequences. Comparing these datasets (Fig. 1A) the BOLD data represented 245 of the 278 UK bee species, but comprised only 14 sequences (6 species) from specimens collected in the UK. The 355 new sequences add 10 UK species (15 sequences) not represented in the BOLD dataset, and novel haplotypes for 107 further species (201 sequences). The final reference set included 255 bee species (92.4% of 278 species known from the UK). The missing species are either extinct (6 species), rarely introduced by accident (1 species, *Heriades rubicola*), only found in the Channel Islands (1 species, *Andrena agilissima*), listed as endangered (RDB3-RDB1) (8 species), or rare and localised (5 species), while 2 species were only recently added (Cross & Notton 2017; Notton *et al*. 2016). When considering each of the six families separately, the greatest number of species missing from the database was in Andrenidae (9 of 69 species), followed by Apidae (4 of 76) and Halictidae (4 of 62).

Genetic variation within morphologically identified Linnaean species ranged from 0% to 5.9% (mean 0.31%, standard error ±0.04%), and interspecific variation ranged from 0% to 24.9% (mean 6.7% ±0.08%). We found that 242 (94.9%) of cox1-based sequence clusters at 97% similarity were an exact match of the Linnaean species identifications (Supplementary Figure S1). Inconsistency with the morphological species definitions were limited to five genera, *Andrena, Bombus, Colletes, Lasioglossum* and *Nomada*.

De novo species delimitation from the DNA sequences using the GMYC method were based on phylogenetic trees generated for each genus (see Fig. 2 for the genus *Nomada*). In most cases of incongruence, the GMYC either split (42 cases) or lumped (14 cases) an existing nominal species, but in rare cases the patterns of splitting and lumping were more complex (Fig. 3). The GMYC species largely agreed with the distance-based BIN network method in the extent to which nominal species were split and lumped (Fig. 2, 3). Inconsistencies of Linnaean and cox1-based entities were mainly due to groups of close relatives with challenging morphological identifications. Subsets of species not monophyletic with respect to each other (a requirement of the GMYC method) included: *Andrena bimaculata - A. tibialis, A. clarkella - A. lapponica - A. helvola - A. varians*, the recently subdivided *Colletes succinctus* species group *(C. halophilus - C. hederae - C. succinctus*] (Kuhlmann et al. 2007), suspected geographically confined species among the *Dasypoda hirtipes* group (Schmidt *et al*. 2015), and variation among *Lasioglossum rufitarse, Nomada flava - N. leucophthalma - N. panzeri*, and *N. goodeniana - N. succincta* clusters.

**Figure 2.**
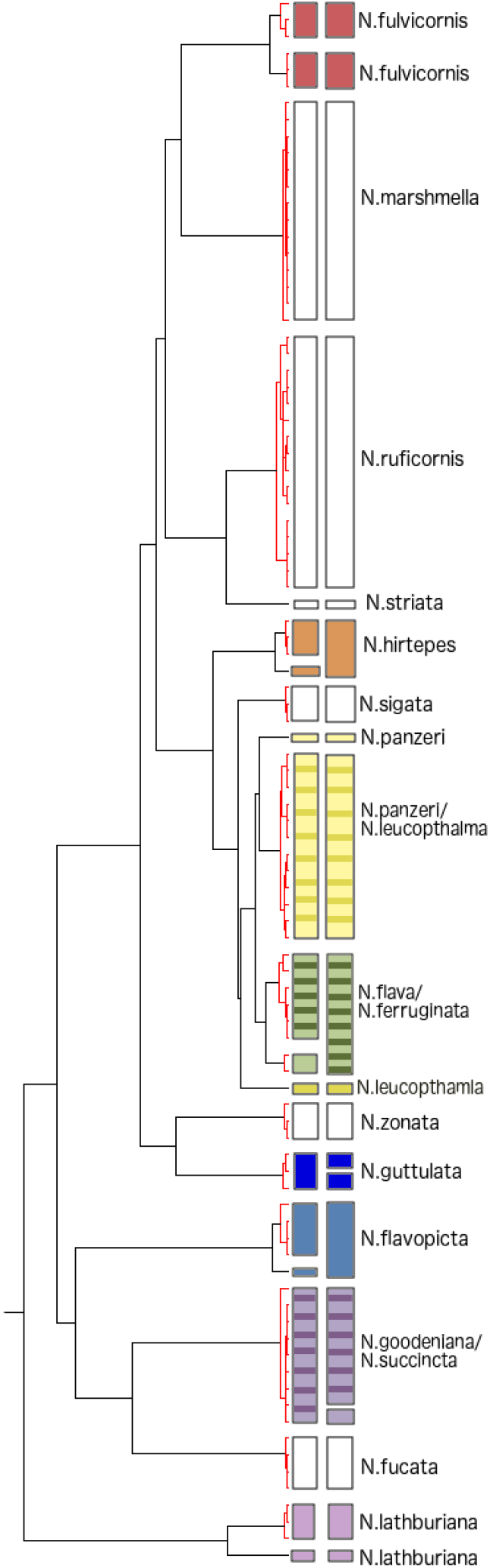
GMYC and BOLD analysis of a subset of the genus *Nomada*. The first column of boxes demonstrates the GMYC species, and the second column of boxes the BOLD bins. Boxes with no fill show species which are not split or lumped with other species in both the GMYC and BOLD analysis. Each colour represents a different species which is either split, lumped or both in either the GMYC or BOLD analysis, or in both. The species names are shown on the tree.

**Fig. 3.**
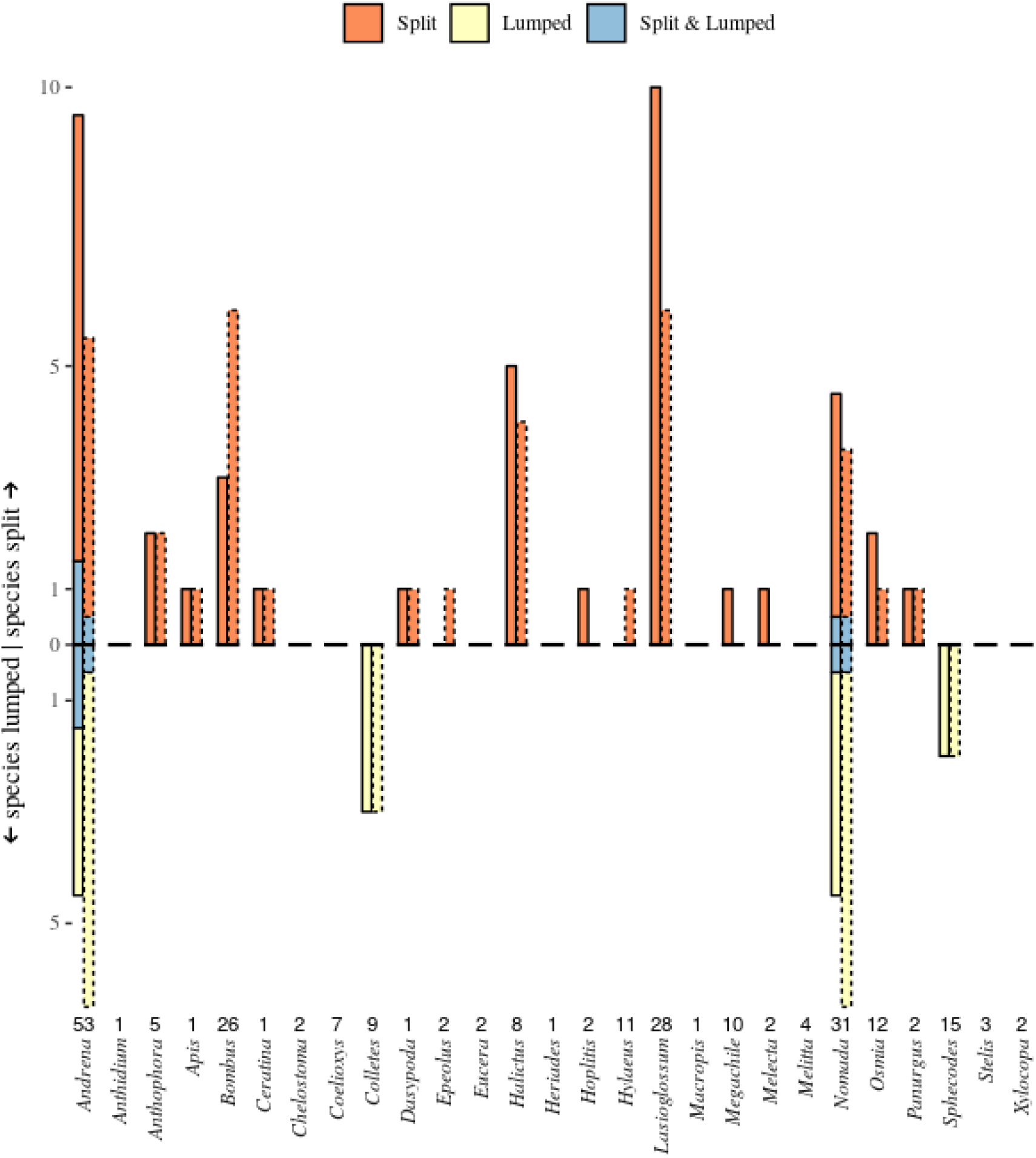
Congruence of species delimitation with assignment to Linnaean species, comparing the Generalised Mixed Yule Coalescent model (GMYC) (solid lines) and BOLD BIN assignments (stipled lines). Each genus is assessed separately. The number of incongruent clusters are shown, either splitting the morphospecies (orange), lump the morphospecies (yellow), or both split the morphospecies and lump those sequences with other morphospecies (blue). The total number of species in each genus is given above the genus name. Note that for many genera the morphospecies assignments were perfectly congruent with either DNA-based methods (no bars).

### Testing HTS data against the reference library

Illumina reads generated for 762 bee specimens resulted in an average of 5851 *cox1* sequences per specimen (amplicon pool) after read merging and stringent quality filtering. Three methods were used to designate a HT barcode from these sequences for each specimen (see Materials and Methods). The NAPselect method, which validates barcode selection by statistical significance of read abundance and taxonomy, obtained a barcode for 749 individuals, failing to do so for 13 samples that did not produce a dominant (hymenopteran) read, while the OTU clustering and most-frequent read method produced only 559 and 584 HT barcodes, respectively (Table 1A). Out of the barcodes chosen by NAPselect, 734 (99.7%) produced a match to sequences in the BEEEE reference data (Table 1A), confirming these sequences correspond to the target specimens. This proportion of hits to the reference set was equally near 100% with the other two methods, but because these produced HT barcodes for fewer specimens, they resulted in approx. 25% fewer specimen identifications. Almost all identifications were to the species level, while between 0 and 3 individuals produced hits to reference sequences identified only to genus (Table 1A). Across all samples, a total of 154 unique species identifications against the BEEEE reference set were obtained.

**Table 1.** The recovery success of different methods of barcode selection and the rate of accurate identification of barcodes against the BEEE reference set. Table 1A. The number of sequences obtained, the number of matches to a sequence in the reference collection and proportion of those that produce a species or genus level identification, respectively. Table 1B. The accuracy of identification, relative to the morphological identification of the specimen, at different levels of morphological or molecular identification. Note that the NAPselect method returned the highest absolute number of correct identifications.

|  |  |  | Most frequent OTU | Most frequent read | NAPselect |
| --- | --- | --- | --- | --- | --- |
| A |  |  |  |  |  |
| Of 762 total specimens: | Specimens with sequences |  | 762 | 762 | 749 |
|  | Specimens with sequences matching reference dataset |  | 559 | 584 | 734 |
| Of 761 specimens with species-level morphological identifications and 1 with genus-level identification: | Sequences with species-level molecular identification |  | 556 (99.5%) | 584 (100%) | 732 (99.7%) |
|  | Sequences with only genus-level molecular identification |  | 3 (0.5%) | 0 (0%) | 2 (0.3%) |
|  |  |  | Most frequent OTU | Most frequent read | NAPselect sequence |
| B |  |  |  |  |  |
| Morphological ID level | Molecular ID level |  |  |  |  |
| Species | Species | Total comparisons | 555 | 583 | 731 |
|  |  | Species-level correct | 471 (84.9%) | 506 (86.8%) | 611 (83.6%) |
|  |  | Genus-level correct | 528 (95.1%) | 565 (96.9%) | 707 (96.7%) |
| Species | Genus | Total comparisons | 3 | 0 | 2 |
|  |  | Genus-level correct | 3 (100%) |  | 2 (100%) |
| Genus | Species | Total comparisons | 1 | 1 | 1 |
|  |  | Genus-level correct | 1 (100%) | 1 (100%) | 1 (100%) |

Congruence of molecular identifications with the morphological identifications of the source specimens was high at genus level with 95-96%, but only 83-86% of specimens were identified as the same species with both data types (Table 1B). However, as NAPselect designated a considerably larger proportion of barcodes to species level, the absolute number of correct species identifications using this method was the highest, at 611 specimens out of the 762 sequenced (707 correct at genus level). The proportion of successful molecular identification was compared between different genera to examine whether there was a taxonomic bias in identification success. The success rate of molecular identification differed among genera (Figure 4), although this tends to be correlated with the number of species/sequences in a genus. Species-rich genera that produced markedly more successful identifications include *Andrena* and *Bombus*, whereas *Colletes* showed low success even using NAPselect (as expected because some species were inseparable by DNA; see above).

We investigated whether the lumping and splitting observed in the reference dataset was a driver of molecular misidentification by examining the proportion of correct and incorrect matches against species that were lumped and/or split in the GMYC analysis. Of the 734 HT barcode sequences generated by NAPselect that had a BLAST match to a BEEEE reference sequence, 17 were to a species that was lumped, 178 to a species that was split and 1 to a species that was both lumped and split. The proportion of correct species and genus level matches for these sets of HT barcodes was very similar to the overall rate: 76.5% of matches to lumped species and 88.2% of matches to split species were correct at the species level (94.1% and 98.8% at the genus level), and the single HT barcode matching a lumped *and* split species was correct at the species level as well.

### Exploration of concomitant DNA in the testing dataset

When OTU clustering was carried out on the entire data set combining the reads from all 762 samples, USEARCH within NAPcluster generated 498 OTUs, of which 263 were identified as Apiformes using BLAST/MEGAN. Out of these, the tree-based assessment of potential pseudogenes identified 72 OTUs as likely NUMTs. In addition, several OTUs were reclassified as Diptera in the phylogenetic tree used for pseudogene filtering. The final count of *bona fide* OTUs identified as Apiformes was 180, of which 170 had hits to the BEEEE reference library. Apiformes thus dominated the set of OTUs, but the dataset also included 235 OTUs from across the eukaryotes, including Diptera (48 OTUs), Coleoptera (6 OTUs), and various other insects (22 OTUs). The Diptera included several species of hoverflies (Syrphidae), which were present in the traps and were processed alongside the bees, but are not discussed here. Five of the six Coleoptera OTUs were identified as common flower visitors, including three species of *Cantharis* Soldier Beetles (Cantharidae), Malachite Beetles (Malachiidae) and a Pollen Beetle (Nitidulidae), in addition to *Zophobas atratus*, a non-native species of Darkling Beetle (Tenebrionidae). There were also OTUs from organisms that associate directly with bees such as Acari (mites) and *Wolbachia* (alphaproteobacteria), as well as several flowering plants and numerous fungi and oomycetes. The Acari comprised four OTUs, of which one was identified to species, *Locustacarus buchneri*, a known tracheal parasite of bumblebees, while the others were identified only as members of the Sarcoptiformes, Crotonioidea and Parasitiformes. Finally, *Homo sapiens* DNA was detected in numerous samples.

Quantitative comparisons of OTU distributions across samples were conducted after rarefaction, which removed 46 samples with fewer than 400 reads, losing 15 OTUs. Rarefied samples had between 1 and 26 OTUs, with a mean of 5.9 (SD = 3.9). The majority of samples had secondary OTUs beyond the specimen barcode sequence. Secondary OTUs contributed an average of 25.7% (SD = 33%) of the samples reads; in 238 cases, secondary OTUs contributed over 50% of the reads. The taxonomic composition of secondary OTUs (Fig. 5) showed that most samples had at least one other bee OTU, sometimes as many as 8, out of the 132 total Apiformes OTUs that were recognised as secondary OTU at least once (48 OTUs of Apiformes were only recovered as the primary OTU). Beyond Hymenoptera, high OTU numbers were contributed by Diptera, with up to 10 OTUs in a single sample, and fungi (maximum 13 OTUs in one sample and a total of 40 across all samples) (Fig. 5). However, no higher taxon was found consistently across all samples apart from the bees.

The high incidence of NAPselect barcode sequences (i.e. Apiformes) occurring as secondary OTUs raised the question about the origin of these non-target specimens in the barcoding mix. Potential sources of DNA may be carry-over from the traps, mixing of specimens during handling for taxonomic identification, and errors in various DNA laboratory procedures. In general, the level of direct contamination with DNA sequences that were the primary OTU in another sample was low, but significantly greater than zero for most sources and source combinations (Supplementary Table SI). Altogether, 132 of the 180 Apiformes OTUs were recognised as secondary OTUs in at least one sample, and 110 of these match to one of the barcode sequences from other wells. Compared with the control, i.e. the background level of cross-contamination from any source, there was a significant increase in contamination rate for within-plate contamination and within-plate and trap contamination (Fig. 6, Supplementary Table S1), indicating that the greatest rate of contamination may have been at the level of library construction, i.e. from mixing among the 96 illumina tags. The level of cross-contamination was much lower for those samples in the same well, i.e. the combination of 13 different products from the primary PCRs conducted with a different primer tag each.

The low level of contamination was reflected in the pattern of cross-contamination of individual species. OTUs identified to 23 different species were each found as secondary OTUs in at least one other sample of a different species from the same trap. The most frequent of these was *Lasioglossum malachurum*, of which there were 37 specimens in the study from 21 traps. We HT barcoded 63 specimens of other species from these 21 traps, and *L. malachurum* was found in 13 of these, a rate of 20%. At trap level, the average rate for the 23 species was 7.6% (SD = 4.5). The same analysis for plates and wells showed that *Lasioglossum calceatum* was the most common cross-contaminator here, being found in 7% of samples of other species sharing a plate (PCR tag) with specimens of *L. calceatum*, and 5% of samples of other species sharing a well (MID) with *L. calceatum*. 45 species cross-contaminated within plates, with a mean rate of 2.2% (SD = 1,7), and 13 species cross-contaminted within wells (mean = 2.5%, SD = 1.1).

## DISCUSSION

### The reference database

Cost-effective species-level identifications of bees and other insect pollinators are required to provide robust evidence for population changes and to inform land use management and conservation (Gill *et al*. 2016). We conducted this analysis in two stages, by first building the reference database using conventional sequencing technology, which was then trialled for species identification using high-throughput sequencing of samples from a proof-of-concept monitoring scheme. The combined effort of new sampling and sequencing, together with barcode data already in the BOLD database, resulted in a virtually complete set of the UK bees, with only a few rare or presumed extinct species missing. Furthermore, we expanded existing references by generating novel sequences from UK populations of widespread species. The *cox1* barcode delimited 94.9% of species in the reference database as separate entities, showing that for almost all bee species in the UK this set is sufficiently discriminatory. In the remaining cases the molecular analysis lumped the Linnaean species, as evident in the *de novo* species delimitation using the GMYC method, while an even greater proportion were shown to be split into additional GMYC groups which, however, were not incongruent with the Linnaean species.

The overall reference database comprises a mixture of UK and non-UK sequences, as many species are more widely distributed in Europe and North America from which many barcodes were obtained, and the species discrimination may be even clearer if performed with UK samples only, as a local subset of intra-specific variation exacerbates the species-level differences (Bergsten *et al*. 2012). Importantly, the high congruence of molecular groups with the Linnaean species also shows that the mitochondrial ‘gene trees’ are a good reflection of the species-level entities, as both morphological diagnostics and mitochondrial markers corroborate the species hypotheses (DeSalle *et al*. 2005), and thus the use of multiple markers for species delimitation is generally not required.

Finally, congruence with the BOLD database also suggests that the identifications have been correct, in some cases after secondary inspection of specimens.

The molecular data failed to separate a small number of species in four of the 27 genera studied (“lumped” in Fig. 3). In some instances, such as the *Colletes succinctus* species group, morphological identification of three named species is reliable, if challenging, now that there is a key covering all UK species (Falk & Lewington 2015), and there are biological and distributional differences. *Cox1* sequences are not sufficient to delineate these species (Kuhlmann *et al*., 2007), and morphotaxonomy remains the most reliable method for this species group. Similarly, the separation of the *Nomada goodeniana-succincta* group relies on subtle colour variants (Falk & Lewington 2015) and cryptic species are likely to exist. Additional genetic markers may be useful; e.g. the three recognised *Colletes* species lumped in *cox1* exhibit fixed differences in EF-1a and ITS (Kuhlmann 2007). Vice versa, divergent *cox1* entities (splitting) may indicate the existence of hitherto unrecognised species. For example, a divergent haplotype in *Dasypoda hirtipes* has now been associated with a morphologically differentiated, eastern European species that is not part of the UK fauna (Schmidt *et al*. 2015). We have already curated the *cox1* database extensively, in particular to remove morphological identification errors (Supplementary Text), but the new clusters may lead to the discovery of separate entities within the Linnaean species and may provide fertile ground for future morphological work. Since DNA extraction destroyed only one leg, morphological vouchers can be re-examined, an important process in refining the reference database.

### Generating high throughput barcodes

The newly created *cox1* database was then used to identify species from a survey of pan traps using high-throughput barcoding (“HT barcoding”). The methodology has great potential for sequencing mixed samples (metabarcoding) but was here applied on individual specimens to test the efficacy of this approach and our ability to confidently recover a sequence for the target specimen. We employed three methods for designating this sequence from a pool of anonymous amplicons. The most intuitive approach was to undertake a standard metabarcoding analysis using the *USEARCH* pipeline to designate the centroid sequence of the most highly represented OTU in each sample as the HT barcode sequence. However, the sequence obtained with this method did not produce a BLAST hit to the reference database in 27% of cases. An alternative method was to simply select the most frequent unique sequence in the amplicon pool, analogous to the sequence that would be generated by Sanger sequencing. However, while this method also designates a barcode for every sample, these sequences are only marginally more likely to find a match to the reference database (23% did not produce a BLAST hit).

The third method, implemented in the NAPselect script, also selects the top-abundant read, but requires that this read matches a specific taxonomic group (in this case, Hymenoptera), and that the read frequency is significantly greater than frequencies of other reads. If these conditions are not met, NAPselect then discards the top read and checks other reads according to descending abundance. This pipeline did not output a sequence for 13 specimens, disregarding samples with low read numbers or low differentiation among other abundant reads. However, the majority of the remaining sequences matched the reference database, and only 3.7% of specimens did not produce a sequence with a BLAST hit - a substantial improvement over the other methods (Table 1). The key improvement introduced by this script probably was that NAPselect conducts BLAST searches against GenBank and assesses the taxonomy of the hits, which was specified to allow BLAST errors. This method is clearly very effective, with error rates determined largely by sequencing depth issues rather than an inability to select the correct sequence.

### Exploration of concomitant DNA in the testing dataset

Unlike standard metabarcoding conducted on mixed samples, the current analysis permits a precise determination of amplicons derived from single specimens. A surprising finding was the high proportion of reads attributable to secondary OTUs, and their taxonomic diversity. The specimens from the monitoring program were not substantially different from those used in Sanger sequencing to build the reference database (in some cases used for both purposes), which produced clean base calls consistent with a single predominant PCR product. However, the primers for illumina sequencing were designed for broad amplification of arthropods (Arribas *et al*. 2016) and probably have a wider taxonomic amplitude than the Hymenoptera-specific primers used to amplify the standard ‘barcode’ region. Besides co-amplification of a broader range of associated species, this may also increase the potential for sequencing of pseudogenes. Out of 509 OTUs recovered from all samples combined, 263 were identified as Apiformes initially, which greatly exceeds the number of species expected in this survey. Pseudogenes diverge without the constraints of coding regions and thus can be partially eliminated based on length differences. For example, a preliminary analysis that did not constrain the read filtering to the target length of 418 bps obtained six additional OTUs assigned to humans, all of which were confidently identified as known mitochondrial pseudogenes. However, filtering the reads to a fixed length could not avoid this problem sufficiently. We therefore implemented a further filter based on the distribution of low-abundance OTUs that are co-distributed with the true mitochondrial copies. We only removed OTUs that form a clade with the presumed true copy (close matches to the reference database), under the assumption that nuclear pseudogenes are of limited evolutionary persistence before they diverge too far from the mitochondrial ancestor and no longer are captured by the PCR primer. Based on these criteria a total of 72 OTUs were identified as mitochondrial pseudogenes. This method (and the removal of several other OTUs whose incorrect assignment was revealed with the phylogeny) reduced the total number of Apiformes OTU to 180, which is closer to the 154 species identified morphologically, in particular if OTU splitting (Fig. 4) is taken into account. The procedure for identifying these likely pseudogene OTUs is a novel step in the metabarcoding filtering process which, to our knowledge, has been implemented here for the first time. However, it is dependent on “true” OTUs being identified by a reference collection and that there exists a high level of read variation between the set of target *cox1* OTUs and their putative pseudogenes – both situations that are common in HT barcoding studies but less so in some metabarcoding. Here, it proved to be a critical step preventing the overestimate of species richness frequently seen in metabarcoding studies.

**Figure 4:**
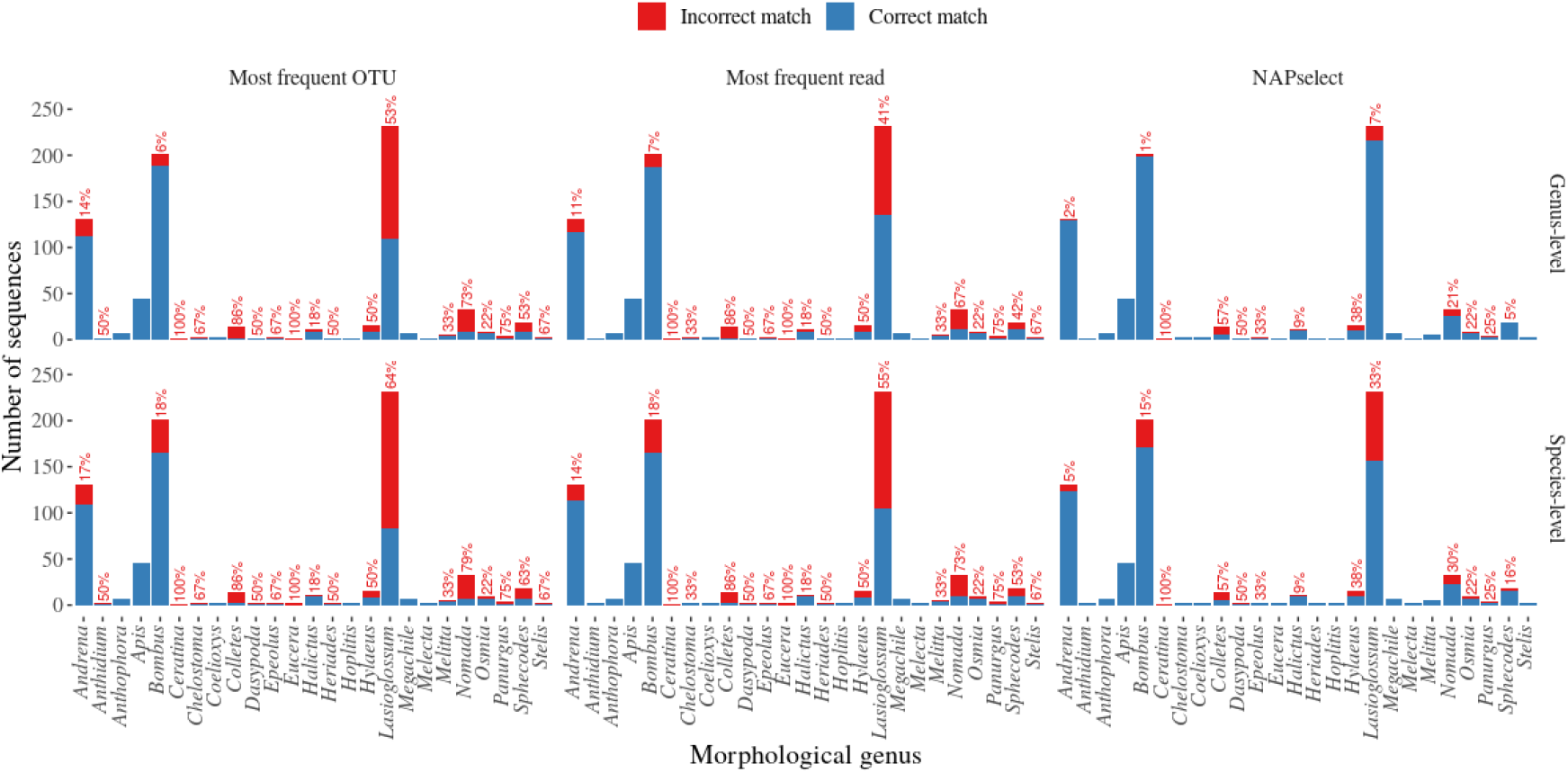
The proportion of molecular identification failure for different morphological species across genera. For each morphological species, we calculated the proportion of specimens for which the designated barcode failed to be correctly identified using the reference database. These values are presented here, grouped by genus and the three different barcode designation methods. Values along the x-axis show the number of morphological species in the genus, and the total number of specimens.

**Figure 5:**
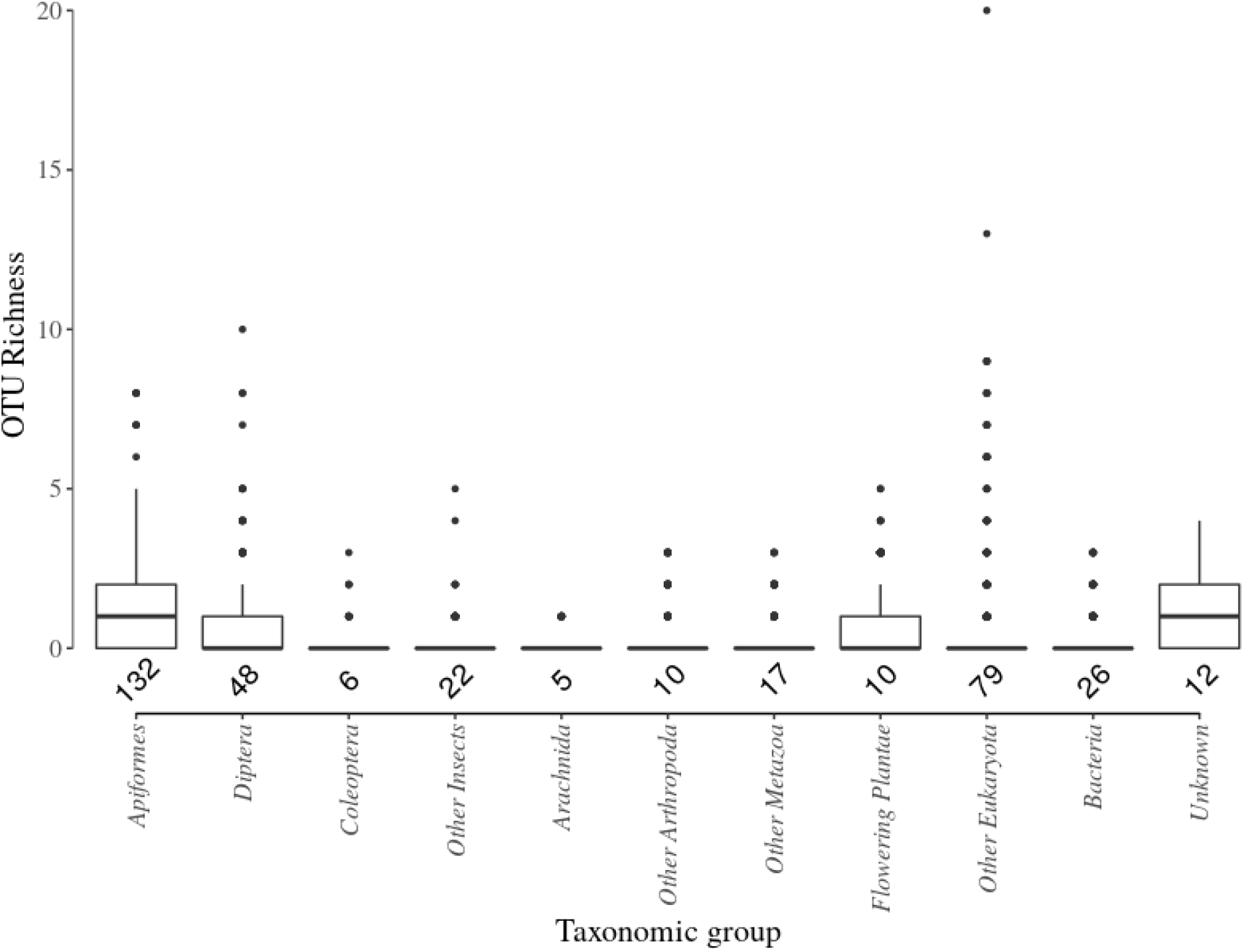
The taxonomic composition of secondary OTUs in NGS barcoding of bees. Boxplot shows the average number of secondary OTUs within major taxonomic groups found in each sample. Values below boxes give the total number of OTUs for that taxon found across the dataset.

**Figure 6:**
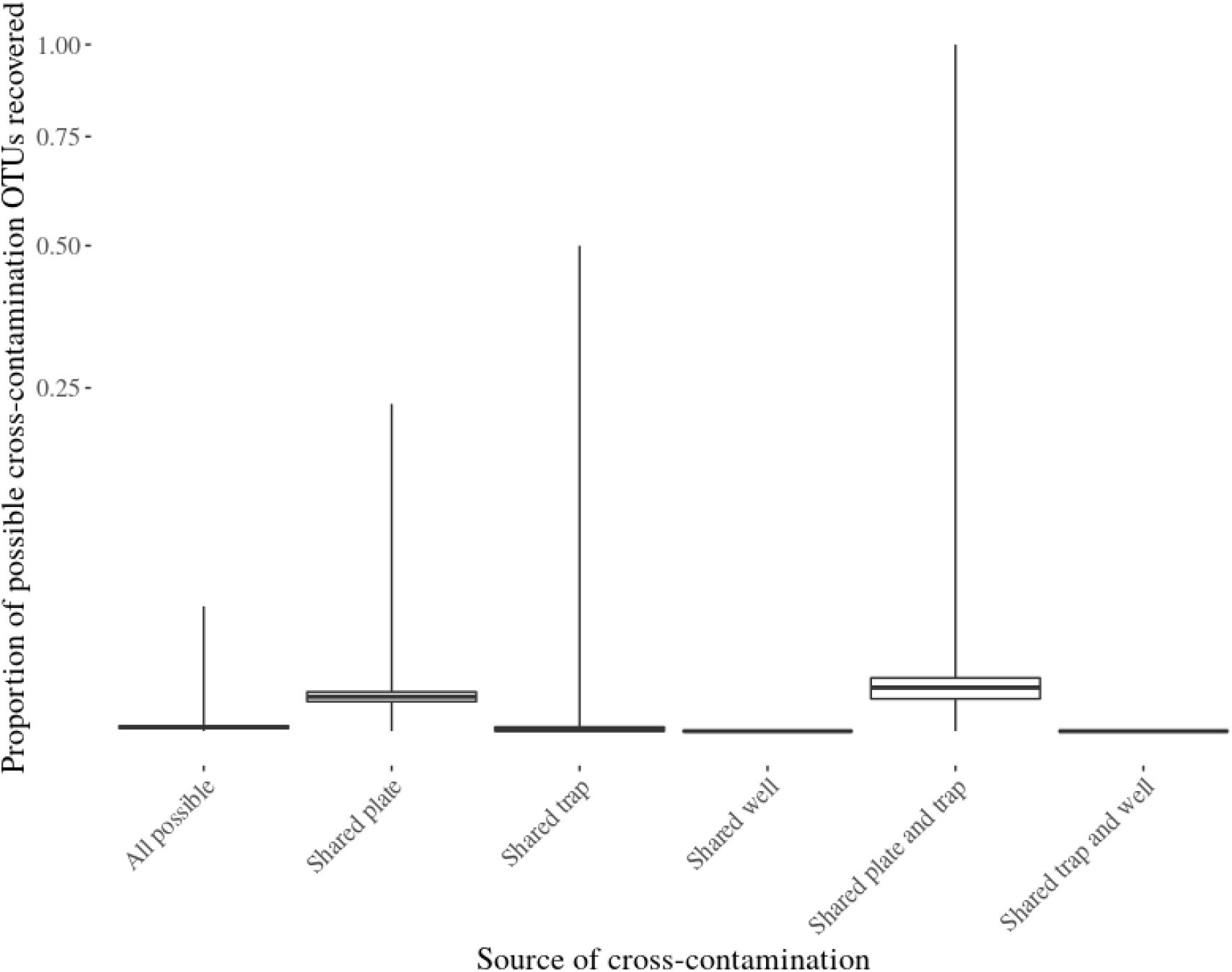
Boxplot showing the rate of recovery of all possible cross-contamination OTUs from different sources of cross contamination. The rate of shared OTU recovery is significantly higher when considering samples from the same plate and same plate and trap compared with a background rate of cross contamination (all possible).

Other secondary OTUs were assignable to a wide range of distantly related taxa, including highly plausible associates of pollinator communities, which suggests carry-over of DNA with the target specimens. Extraneous insect species in the sequencing mixture mostly consisted of other known pollinators attracted to flowers (and pan traps), including various Coleoptera and Diptera. Consistent with the detection of pollen beetles (*Meligethes*), numerous specimens were observed in the pan traps. Species of Diptera included the wheat stem borer *Cephus pygmeus*, a flower visitor whose larvae feed in the stems of cereal crops and wild grasses (Poaceae), and *Sarcophaga* sp. (flesh flies) that are carrion feeders or parasitoids of other invertebrates. The greatest proportion were hoverflies (Syrphidae); these were widely present in the traps and were processed in a parallel study in the same sequencing run and thus additionally exposed to the risk of laboratory contamination as well as trap contamination. Other sequencing records were consistent with internal parasites, including a species of tracheal mite, *Locustacarus buchneri*, known to be associated with bumble bees (*Bombus* sp.), and numerous bacterial sequences. OTUs belonging to Angiospermae suggest the types of flowering plants pollinators visited, including *Caryophyllales* sp., *Cichorieae* sp., *Geraniaceae* sp. and Lamiids (a large clade of flowering plants that includes many species present in meadows). In addition, widely observed ‘unknown’ OTUs to which MEGAN could not confidently assign an identity may be members of taxa that were poorly represented in GenBank, or they may be chimeras or sequencing errors that escaped filtering. Yet, most secondary OTUs are plausible as true associates of the target specimens and the wider pollinator community. Thus, associated DNA can be used to detect local community composition and ecological associations, including parasites, symbionts and diet of the target.

Cross contamination in the traps may also explain the large number of secondary OTUs assigned to Apiformes (beyond the pseudogenes). The potential for DNA mixing was further increased as specimens from the same pan trap were stored together prior to morphological identification and DNA extraction. However, we find that the greatest rate of contamination may have been within a single plate, i.e. between samples with the same primer index but different library indices, which could be either due to physical mixing in the laboratory, tag-jumping (in the library indices, not the PCR tags), or errors in index sequencing. Trap-level contamination may add to the problem, as the combined model (plate x trap) shows only marginally higher levels of contamination (Supplementary Table S1). Because the contamination within the wells was much lower, we conclude that the primary PCR using 13 different primer tags before being combined in a single Nextera XT library was not greatly affected by these problems, indicating that our approach of using the same unique primer tags on forward and reverse strands can largely eliminate the problem of misassignment of PCR fragments. In addition, some types of contamination were less likely to be introduced during molecular lab processing, given the precautions with specimen handling and the strict protocols of the sequencing facility, in particular regarding the widely found human DNA, present in virtually every one of the specimens. As scientists using morphological and molecular methods work together, greater awareness of these issues is needed and the steps to avoid DNA contamination should be understood and implemented, such as the use of clean pans, bee nets and storage bottles, and use of latex gloves for specimen handling during morphological identification.

### Conclusions

High-throughput sequencing can greatly change the approach to monitoring of pollinators, through mass identification of sequence reads against reference databases verified by taxonomic specialists. In this proof-of-concept study we used individuals, rather than bulk samples, to study the outcome of metabarcoding in greater detail. We first established the power of the *cox1* marker for species discrimination, which only left about 5% of UK species without a precise identification at species level. The subsequent utilization of the database for UK bees monitoring shows high consistency with morphological identifications conducted in parallel. However, the deep sequencing of single specimens also revealed the various pitfalls of metabarcoding. We detected surprisingly high levels of apparent mixing with other specimens from the same and other traps. In addition, we found numerous OTUs apparently contributed by pseudogenes, which greatly inflate estimates of the total species diversity; they can be filtered out efficiently as their distribution ‘trails’ the actual mitochondrial copies, which should be a routine part of the read filtering procedure. Lastly, the widely used OTU clustering may not produce the most accurate species detection, as shown by a comparison of OTU analyses against the most abundant read in each sample (after adequate taxonomic and numerical filtering), which revealed a full identification of the target specimen in approximately 25% more samples. Yet, applied under stringent quality filtering, it is possible to use high-throughput sequence data at the read level, i.e. to establish genotypic variation or for assignment to particular subgroups within the Linnaean species, and thus use them in the same way as data from Sanger sequencing, but scaled up by orders of magnitude. The method thus greatly increases the accuracy and speed of taxonomic identification in pollinator monitoring, at reduced cost, while also providing further information on species interactions and ecosystem composition through the secondary OTUs. The bioinformatics methodology and comprehensive barcode database can now be rolled out for the study of much larger number of specimens typically obtained by passive pan traps and can be extended to studies of pollinators in other parts of the world.

## ACKNOWLEDGEMENTS

This work was funded by the UK Department of Environment, Forestry and Rural Affairs (Defra) of the UK (contract PH0521), with in-kind contributions from the NHMUK, and a fellowship of the NERC Science Solutions for a Changing Planet Doctoral Training Programme at Imperial College (to HN). The bioinformatics pipelines were developed under the iBioGen project funded by the European Commission. The NPPMF pilot pan trapping study was jointly funded by Defra and Scottish Government under project WC1101. We acknowledge the Borough of Lewisham (Blackheath), Bristol City Council (The Downs, Troopers Hill), Conservators of Wimbledon and Putney Commons, Land Trust (Greenwich Peninsula Ecology Park), National Trust (Bookham Common, Leigh Woods), Natural England (Hartslock SSSI), and Royal Borough of Greenwich (Blackheath) for permission for DGN to collect bees. Jackie Mackenzie-Dodds (NHMUK) and NPPMF staff are thanked for making the NPPMF collection available. Martin Harvey, Stuart Roberts and Ivan Wright conducted the morphological identifications of bees sampled in the NPPMF pilot study.

## Data accessibility

Sequence data available at BOLD under the BEEEE label. Perl scripts used for the sequence clustering and barcode selection are available at https://github.com/tjcreedy/NAPtime.

## Author contributions

CQT, HN and APV designed the study; CQT and HN generated molecular data; TJC, CQT, KQC and HN performed data analysis. TJC developed bioinformatics tools. CC, KC and RO collected specimens and co-ordinated morphological identifications. DGN collected specimens, identified, documented, sampled them, preserved morphological vouchers, and verified identification. PA and CA designed analytical pipelines and provided advice on project design and analysis. CQT, HN, TJC, KQC and APV wrote an initial draft of the manuscript. All authors contributed to the writing of the final draft.

